# Low competitive status elicits aggression in healthy young men: behavioral and neural evidence

**DOI:** 10.1101/2020.06.20.162586

**Authors:** Macià Buades-Rotger, Martin Göttlich, Ronja Weiblen, Pauline Petereit, Thomas Scheidt, Brian G. Keevil, Ulrike M. Krämer

## Abstract

Winners are commonly assumed to compete more aggressively than losers. Here, we find overwhelming evidence for the opposite. We first demonstrate that low-ranking teams commit more fouls than they receive in top-tier soccer, ice hockey, and basketball men’s leagues. We replicate this effect in the laboratory, showing that male participants deliver louder sound blasts to a rival when placed in a low-status position. Using neuroimaging, we characterize brain activity patterns that encode competitive status as well as those that facilitate status-dependent aggression in healthy young men. These analyses reveal three key findings. First, anterior hippocampus and striatum contain multivariate representations of competitive status. Second, interindividual differences in status-dependent aggression are linked with a sharper status differentiation in the striatum and with greater reactivity to status-enhancing victories in the dorsal anterior cingulate cortex. Third, activity in ventromedial, ventrolateral, and dorsolateral prefrontal cortex is associated with trial-wise increases in status-dependent aggressive behavior. Taken together, our results run counter to narratives glorifying aggression in competitive situations. Rather, we show that those in the lower ranks of skill-based hierarchies are more likely to behave aggressively and identify the potential neural basis of this phenomenon.

## Introduction

Winners are often taken to be aggressive, not only in sportive environments (Chow *et al.*, 2009; Trebicky *et al.*, 2013) but in society at large (Waasdorp *et al.*, 2013; Laustsen and Petersen, 2017; Cheng, 2020; Weick, 2020). Indeed, adjectives such as “aggressive” or “dominant” are frequently casted in a positive light in competitive contexts (Næss, 2001; Pappas *et al.*, 2004), and leaders perceived to possess these characteristics are preferred in conflict situations (Kakkar and Sivanathan, 2017). From this perspective, aggression might serve to both achieve and assert a higher social rank (Lee and Yeager, 2020). In agreement with this idea, individuals tend to deliver louder sound blasts to a rival they outperformed than to one who outperformed them (Muller *et al.*, 2012). Contradicting these findings, the lower sense of control and the frustration occasioned by frequent defeats can fuel aggression in subordinate competitors (Przybylski *et al.*, 2014; Oxford *et al.*, 2017; Dowsett and Jackson, 2019). Supporting this notion, a number of studies has shown that individuals are more aggressive towards persons with high status (Davis and Reyna, 2015; Hu *et al.*, 2016; Saalfeld *et al.*, 2018; Kakkar *et al.*, 2019), as low status might impede goal attainment (Berkowitz, 2012), induce stress (Sapolsky, 2004; Sapolsky, 2005) and threaten self-image (Horton and Sedikides, 2009). Furthermore, while accomplished contestants can rely on their ability, less proficient ones might compete more aggressively to outweigh the skill differential (Kirker *et al.*, 2000; Coulomb-Cabagno and Rascle, 2006). Hence, those in the lower ranks of skill-based hierarchies might be more prone to use aggression than those who win consistently. If that were the case, social narratives emphasizing aggressiveness as a requisite for competitive success would be misguided. Here, we tested whether competitive status is related to physical aggression and investigated the neural underpinnings of this association.

Status-based hierarchies order social life and are hence imprinted in human neurobiology (Qu *et al.*, 2017). The rostral aspect of the medial prefrontal cortex (mPFC) has been postulated as a crucial region for inferring a rival‘s dominance during competitive interactions (Ligneul *et al.*, 2016), which might be partially attributable to this region’s role in encoding stable behavioral traits (Hassabis *et al.*, 2013). The amygdala and anterior cingulate cortex (ACC), on the other hand, have been suggested to track moment-to-moment changes in status-based hierarchies (Kumaran *et al.*, 2012; Kumaran *et al.*, 2016). The ventral striatum (VS) and ventromedial prefrontal cortex (vmPFC) are highly responsive to competitive outcomes and might thus also contribute to acquire status representations during skill-based contests (Ligneul *et al.*, 2016). Further, these regions show enhanced reactivity to high- as compared to low-status individuals during both competitive (Zink *et al.*, 2008) and non-competitive tasks (Zerubavel *et al.*, 2015). In addition, there is evidence suggesting that the hippocampus encodes relational knowledge of social hierarchies (Kumaran *et al.*, 2012; Schafer and Schiller, 2018; Park *et al.*, 2020), along with regions involved in social cognition such as the precuneus and the temporoparietal junction (TPJ) (Muscatell *et al.*, 2012; Tavares *et al.*, 2015; Zerubavel *et al.*, 2015). Thus, the current neuroscientific evidence permits to formulate a tentative description of how competitive hierarchies are processed in the human brain. First, regions associated with salience detection (amygdala, ACC) capture dominance signals and status changes during social interactions. Second, regions involved in affective valuation (VS, vmPFC) assess an individual’s current status and facilitate hierarchy learning on the basis of victories and defeats. Finally, the *mentalizing network* (precuneus, TPJ, dorsal and rostral mPFC) and the hippocampus enable the encoding and implementation of status hierarchies during competitive decision-making. The key question that we addressed here is which role these neurocognitive processes play in status-dependent aggression. We reasoned that the neural substrate of status-processing and aggression should show some degree of overlap, be it in subcortical structures assumed to generate aggressive impulses such as the amygdala (da Cunha-Bang *et al.*, 2017; Buades-Rotger and Krämer, 2018), in those linked with retaliation such as the VS (Buades-Rotger, Brunnlieb, *et al.*, 2016; Chester and DeWall, 2016), and/or in areas suggested to regulate aggression such as the vmPFC (Buades-Rotger *et al.*, 2019; Bertsch *et al.*, 2020).

A potentially crucial neuromodulator of status-seeking behavior is the steroid hormone testosterone. Though generally associated with aggression (Geniole *et al.*, 2020), recent evidence indicates that testosterone can flexibly promote prosocial behavior in order to improve one’s social status (Eisenegger *et al.*, 2011; Terburg and van Honk, 2013; Geniole and Carré, 2018). For instance, a study showed that endogenous testosterone was positively associated with generosity in low-status (i.e. junior) rugby players, whereas the relationship was *negative* in high-status (i.e. senior) players (Inoue *et al.*, 2017). Similarly, men with relatively higher testosterone concentrations punished unfair offers more severely, but rewarded fair offers more generously (Dreher *et al.*, 2016). Given that high-testosterone individuals are generally more sensitive to status cues (Josephs *et al.*, 2006; Wagels *et al.*, 2018; Losecaat-Vermeer *et al.*, 2020; Wu *et al.*, 2020; Han *et al.*, 2021), and that competitive situations inherently invoke the use of aggressive strategies (Waddell and Peng, 2014; Dowsett and Jackson, 2019), testosterone might increase competitive aggression by modulating neural reactivity to status signals. Specifically, both preparatory and victory-contingent testosterone surges can induce competitive aggression by activating core neural structures of the threat- and reward-processing systems (i.e., amygdala and VS respectively) (Geniole and Carré, 2018). Importantly, the effect of testosterone on the neural circuitry of competitive aggression is likely to be more pronounced in men (Zilioli and Bird, 2017), who show stronger endocrine reactivity to competition (Geniole *et al.*, 2017) as well as a tighter association between testosterone and aggression (Geniole *et al.*, 2020). The relationship between status-seeking behavior and testosterone might however be moderated by cortisol, a stress-related hormone that is thought to partly suppress testosterone’s effects (Casto and Edwards, 2016; Dekkers *et al.*, 2019). In the present study, we drew on pre-existing sports data, a behavioral study and a neuroimaging study to test a) whether competitive status is linked with aggression in healthy young men, b) whether this is contingent on the neural processing of status signals, and c) whether testosterone amplifies aggressive and/or neural responses to status cues, either by itself or in interaction with cortisol.

## Correlational study

In a first correlational study, we probed whether competitive status was associated with aggressive play across seasons in soccer, basketball and ice hockey. This proof-of-principle approach allowed us to test the link between competitive status and aggression in real-world settings.

### Correlational study: data acquisition

We defined competitive status as a team’s position at the end of the regular season, with higher values indicating a lower standing in the rankings. Aggression was operationalized as the number of fouls (soccer), personal fouls (basketball), or penalty time (ice hockey) incurred relative to received, a measure that accounts for the reciprocal, “tit-for-tat” character of sports aggression. For basketball and ice hockey, we determined the position of teams from different divisions on the basis of their win percentage. We manually extracted data from http://www.footstats.co.uk (soccer), http://www.nba.com (basketball), and http://www.hockey-reference.com (ice hockey) since the last year in which complete, correct, data was available onwards (2000-2008 for European football leagues, 2005 for basketball and hockey). We collapsed these variables separately for the main five European football leagues (England, Spain, Germany, Italy, and France), the North-American National Basketball Association league (NBA), and the North-American National Hockey League (NHL) for a total of 2254 observations (i.e. teams in a given season). By using data aggregated over seasons we minimize the effect of proximal situational influences (home-field advantage, referee biases, etc.) that might induce fluctuations in the number of fouls incurred in single games. The correlation between received and committed fouls or penalties was r_1412_=.74 for football, r_418_=.70 for basketball and r_418_=.93 for ice hockey. This indicates that a) aggressive play follows reciprocal, “give-and-take” dynamics so that teams who commit more fouls also tend to receive more, but b) that the ratio of fouls/penalties committed to received captures some unique variance in teams’ aggressiveness that is not accounted for by the raw amount of fouls or penalty time incurred. This is thus a valid measure to assess the degree to which a team engages in sanctioned aggression.

### Correlational study: data analysis

Because of the ranked nature of the data, we computed Spearman correlation coefficients between position and foul ratio. We also tested whether this relationship held across seasons and leagues and when accounting for the nested data structure. To do so, we ran linear mixed-effects model analyses with position, season, and league as Z-transformed fixed-effect predictors and team as grouping factor. Following best-practice recommendations (Barr *et al.*, 2013), we attempted to maximize the random-effects structure of the model defining by-team random slopes for all fixed effects and random intercepts for all predictors where possible. The most complex model to converge was one with random slopes for season and random intercepts for league, position, and team.

All analyses were performed in R version 3.6.1 running on R Studio 1.1.423. We used the psych package (Revelle, 2017) for Spearman correlations and ggplot2 for plotting (Wickham, 2016). For linear mixed-effects modeling we used the lmerTest package (Kuznetsova *et al.*, 2017). Data and analysis scripts for this study are available via the Open Science Foundation (https://osf.io/2jvx4/).

### Correlational study: results

As shown in Fig. 1, low ranking was consistently associated with foul ratio in each individual league as well as across leagues (mean weighted ρ=.30; Cohen’s d=.63, p<.001). When pooling over all competitions (Fig. 1h), low-ranking teams (Z-transformed position>1) showed on average a 9.5% greater foul ratio (104.68% vs 95.18%) than high-ranking ones (Z-transformed position<−1). Linear mixed-effects models revealed that the relationship between rank and foul ratio persisted (β=.21, t_89_=8.80, p<.001) when controlling for league and season, which had no effect and did not interact with ranking (all p>.124; Table 1).

**Table 1:**
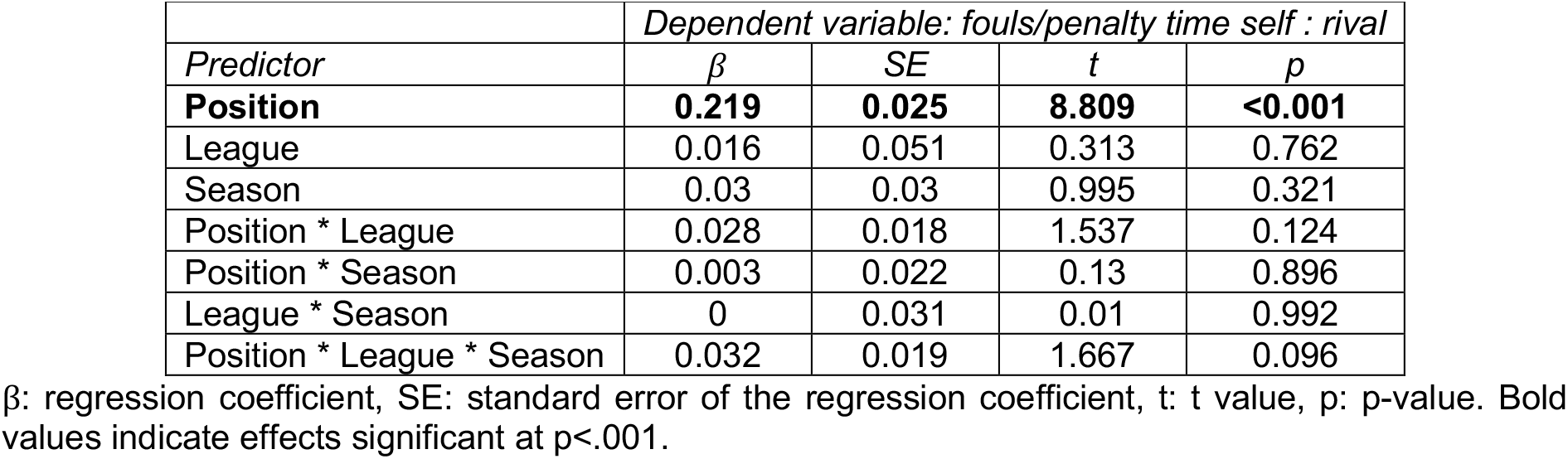
Results of linear-mixed effects model predicting foul/penalty ratio in sports teams (n=2254)

**Fig. 1.**
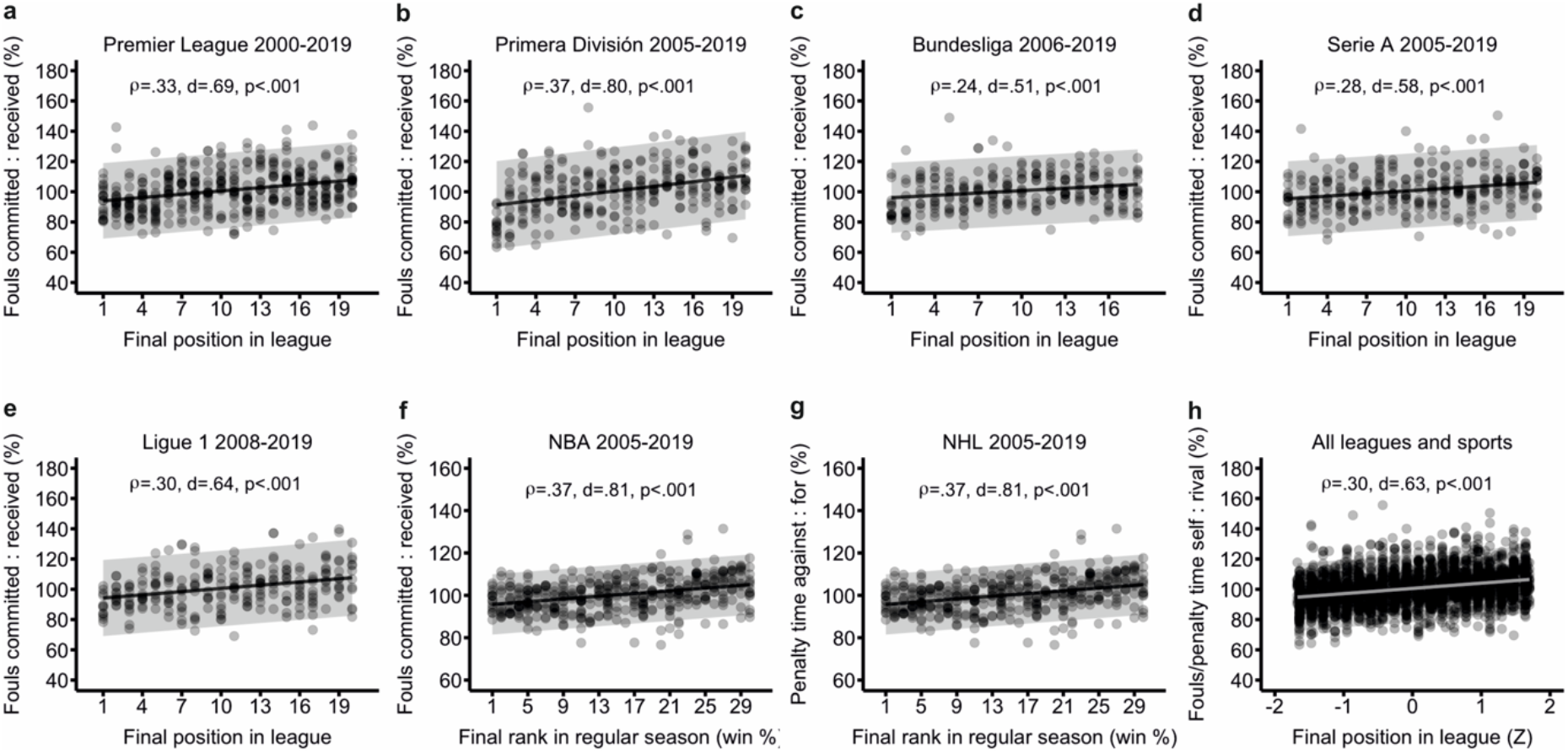
Correlation between competitive status and aggression in elite sports teams. Status was defined as the team’s final position at the end of a season. Aggression was defined as the ratio of fouls (soccer), personal fouls (basketball), or penalty time (ice hockey) incurred relative to received, in %. We collapsed data across seasons. We include best-fit lines and 95% prediction intervals. **a**, English Premier League 2000-2019, **b**, Spanish Primera División 2005-2019, **c**, German Bundesliga 2006-2019, **d**, Italian Serie A 2005-2019, **e**, French Ligue 1 2008-2019, **f**, North-American National Basketball Association League 2005-2019, **g**, North-American National Basketball Association League 2005-2019, **h**, All data points collapsed, with position expressed as Z-scores.

## Behavioral study

Our analysis of sports data provides consistent -albeit purely correlational- evidence for a link between low competitive status and aggression at the level of teams. It is however unclear whether the status-aggression relationship can also be observed in individual competitors. Moreover, fouls in sports serve a predominantly instrumental function. A more stringent test of our main hypothesis would entail manipulating status experimentally during one-to-one contests in which aggression does not strictly fulfil a competitive purpose, i.e. it does not directly impact rivals’ performance. To that end, we ran a pre-registered behavioral study (https://osf.io/q5ge6) in order to probe whether individuals would be more or less aggressive as a function of their competitive status. In order to investigate the effect of status on aggression experimentally, we devised a modified version of the Taylor Aggression Paradigm, a competitive reaction time task in which the winner can choose the volume of a sound blast to punish the loser (Buades-Rotger, Engelke, *et al.*, 2016). Critically, we manipulated competitive status by programming the task so that subjects won more often against one rival than against the other.

### Behavioral study: participants

As this was an exploratory study, we aimed for a sample size of n=30 in order to have 80% power to detect a middle-sized within-subject effect (d=.5) at a conventional two-sided threshold of p<.05. After exclusion of three participants (two saw through the status manipulation and one failed to understand the task, see protocol below), the sample comprised 25 healthy young men (age: 24±3.1; height: 180±5.1 cm; weight: 75.96±9 kg; 23 right-handed, 2 ambidextrous). The study had been approved by the Ethics Committee of the University of Lübeck. Subjects consented to participate and to the anonymized dissemination of the data.

### Behavioral study: protocol and task

Participants were measured in groups of three and believed they would compete against each other, but they actually played against the computer. A male confederate filled in for a participant when an appointment with three subjects was not possible. We first took participants to a computer room in which they provided informed consent and read the instructions together. Computers were separated by screen-walls so that participants could not see each other, and they wore headphones throughout.

We programmed the task so that participants competed in random order, but not more than three times in a row, against each opponent. After eight practice trials, the task started (see Fig. 2a for an outline of the task). Trials began with a decision phase wherein subjects saw the percentage of trials won by their current opponent and by themselves as separate filled bars, i.e. their competitive status. During the decision phase, participants set the volume of a sound blast in a 1-8 scale to be later delivered at their opponent should they win a subsequent reaction time task. The chosen volume was our measure of aggression. In the reaction time task, they had to be quicker than their rival in pressing any button when a target appeared (i.e. a bullseye). We manipulated status so that participants lost more frequently against one rival (66%) than against the other (33%) for a total of 60 trials. To make the setting more believable, subjects were more likely to lose when they were slower than their own median cumulative reaction time, and we interspersed shorter (4s) and longer (12s) inter-trial intervals; during the latter, the opponents allegedly played against each other. Following previous work (Buades-Rotger, Engelke, *et al.*, 2016), we programmed the opponents’ punishment selections to increase progressively over time (i.e. every 20 trials) in order to gradually provoke participants.

**Fig. 2.**
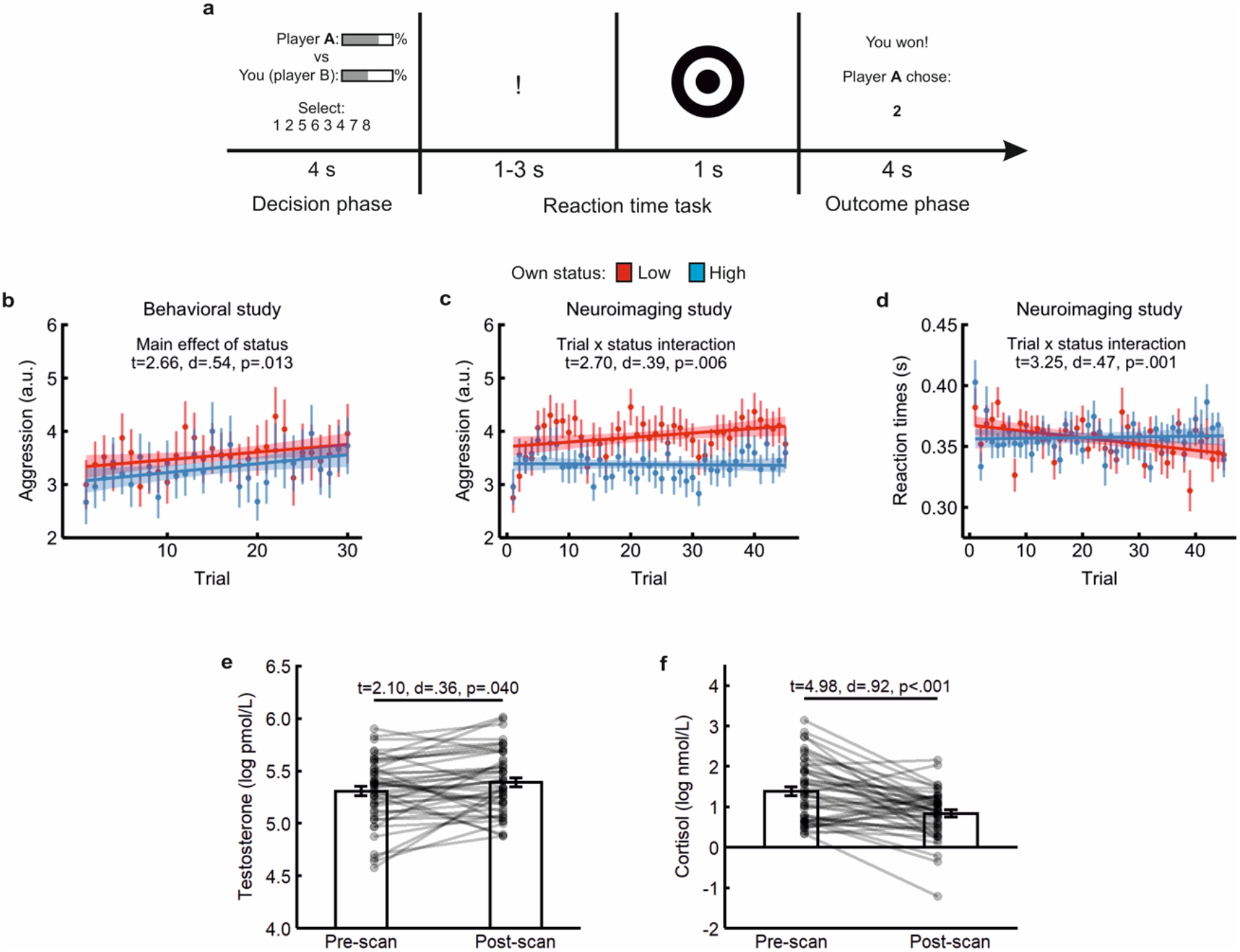
**a**, outline of the task. Subjects saw the % of victories achieved by the opponent and themselves as a filled bar while choosing the volume of a sound blast to be later directed at their opponent (decision phase). Then, they had to be faster than the rival in pressing a button when a jittered bullseye appeared (reaction time task). In the outcome phase they were informed of whether they won or lost and of the rival’s selection. If they lost, they received the sound blast at the end of the trial via headphones. They were consistently worse against one opponent (low-status condition) and better than the other (high-status condition), though both rivals chose equally strong sound blasts on average. **b**, mean punishment selections by trial and participant status in the behavioral study, including best-fit lines and 95% prediction intervals. **c**, mean aggression by trial and participant status in the neuroimaging study, including best-fit lines and 95% prediction intervals. **d**, mean response latency in the reaction time task by trial and participant status in the neuroimaging experiment. **e**, testosterone levels in saliva pre and post-scanning. **f**, cortisol levels in saliva pre- and post-scanning.

After the measurement, participants filled out a manipulation check asking how unpleasant they found the loudest and lowest sound blasts to be with a Likert-type scale ranging from 1 to 8, the perceived average punishment selection of each opponent (also with a Likert-type 1 to 8 scale), and the percentage of trials they believed to have won against each opponent. This questionnaire also probed deception success via three open-end questions: “Did you notice anything special about the behavior of your fellow players?”, “Did you have a particular strategy for the game?”, and “What do you think was investigated in this study?”. We excluded participants if they made explicit mention to the task being preprogrammed (e.g. “The other players were bots”), if they misunderstood the task (e.g. not pressing any button in the reaction time task), or if they evinced an insufficient knowledge of the German language. At the end of the experiment, we debriefed participants regarding the goals and methods of the study.

### Behavioral study: data analysis

Average punishment selections show high internal consistency, load onto a single factor (Chester and Lasko, 2019), and predict real-life aggression (King and Russell, 2019), suggesting that they are a valid measure of aggression and that they adequately summarize participant’s behavior in the task. We followed the pre-registered analysis plan (https://osf.io/q5ge6) and compared mean aggression against the high- vs low-status opponent with a paired t-test in R (version 3.6.1) running on R Studio (version 1.1.423). Data and analysis scripts for this study are freely available via the Open Science Foundation (https://osf.io/2jvx4/).

### Behavioral study: results

Post-experimental manipulation checks revealed that they were aware of winning less often (t_24_=7.55, d=1.51, p<.001) against the better-performing player (34.8% ± 2.1% [mean ± standard error]) than against the worse one (60.5% ± 1.9%). As intended, they did not perceive either rival to be more aggressive than the other (t_24_=1.58, p=.125). Following the pre-registered analysis plan (https://osf.io/q5ge6), we extracted mean punishment selections per condition and compared them with a paired t-test. Subjects selected louder sound blasts in a low- than in a high-status position (t_24_=2.66, d=.54, p=.013; Fig. 2b).

## Neuroimaging study

In a subsequent neuroimaging study, we used a slightly modified version of the same task to scrutinize neural responses to status information during a competitive interaction. Further, we tested whether status-related brain activity was linked with actual aggressive behavior in the task as well as with salivary testosterone and cortisol measured before and after scanning. Unlike the behavioral study, we programmed opponents’ punishments to stay relatively constant over time. We did so in order to disentangle provocation and status effects and to render trials more comparable throughout the task for subsequent averaging of brain activity. We hence set the opponents to select punishments in the middle range (3-6) throughout the task instead of increasing progressively. We also lengthened the task for a total of 90 trials, divided in 3 runs of 30 trials each (15 per opponent) to increase statistical power. In this case, the opponents were always confederates of the experimenters who pretended to play against the participant from computers outside the scanner. To increase the plausibility of this setting, participants played the eight practice trials together with the confederates in the same computer room where the pilot measurements had taken place.

### Neuroimaging study: participants

We determined the sample size on the basis of a previous study (n=39) in which we observed middle-sized correlations (r≈.4) between endogenous testosterone, aggression, and brain activity using a similar task (Buades-Rotger, Engelke, *et al.*, 2016). We thus aimed for a sample size of 50, which yields ~80% power to detect effects of r=.4 or larger with a two-sided threshold of p<.05. After exclusion of six subjects due to non-deception (n=4) and failure to understand the task (n=2), the final sample comprised 47 men aged 24±3.8 (height: 182±6.6 cm; weight: 79.90±11 kg; 37 right-handed, 7 left-handed, 3 ambidextrous) who were reportedly free of current psychiatric, neurological, and endocrine diseases except for one participant with treated hypothyroidism. We included left-handed participants because we did not have specific hypotheses regarding lateralization of the effects and because subjects used both hands to perform the task. This study had also been approved by the Ethics Committee of the University of Lübeck. Subjects provided informed consent for participation and anonymized dissemination of the data. We used the same post-experimental questionnaire as in the behavioral study and we debriefed them regarding the goals and methods of the study. Measurements began between 12 and 15 PM in order to minimize circadian fluctuations in hormonal levels, with the exception of two measurements (17 and 18 PM) that could not be scheduled otherwise.

### Neuroimaging study: acquisition of testosterone and cortisol data

Subjects provided saliva samples in plastic vials (SafeSeal micro tube 2ml from Sarstedt) with the passive drooling technique before and after scanning. We froze the samples at −20°C and shipped them in dry ice to author BGK’s laboratory in Manchester (UK) for analysis once study was completed. Free testosterone and cortisol concentrations were estimated with liquid chromatography tandem mass spectrometry (LC-MS/MS) as described elsewhere (Perogamvros *et al.*, 2009; Keevil *et al.*, 2013). Coefficients of variation (CV) with this technique have been reported to be 5.3% for testosterone and 8.7% for cortisol, whereas mean inter-assay CV were 9% for testosterone and 7.8% for cortisol (Perogamvros *et al.*, 2009; Keevil *et al.*, 2013). The lower limits of quantification (LLOQ) were 5 pmol/L for testosterone and 0.8 nmol/L for cortisol.

### Neuroimaging study: acquisition of fMRI data

We acquired all scans with a 64-channel head-coil mounted on a 3-T Siemens Magnetom Skyra scanner at the Center for Brain, Behavior and Metabolism (CBBM) at the University of Lübeck. For functional volumes we applied a single-shot gradient-recalled echo-planar imaging (GRE-EPI) sequence sensitive to blood oxygen level dependent (BOLD) contrast (TR=1060 ms; TE=30 ms; flip angle=60°; 60 transversal slices; slice thickness 3 mm; in-plane voxel size=3×3 mm^2^; field-of-view [FOV]=210×210 mm^2^, simultaneous multi-slice factor=4; full-brain coverage). We acquired 570 volumes per run for a total of three runs (≈30 minutes). Before and after the task we acquired two magnetic field measurements (2D double-echo gradient-echo sequence; TE_1_=5,17ms; TE_2_=7.63ms; TR=554ms; flip angle=60°; 50 transversal slices; slice thickness 3 mm; in-plane voxel size=3×3 mm^2^; FOV=240×240 mm^2^) for subsequent offline distortion correction as well as two 6.5 minutes eyes-closed resting-state runs not analyzed for the present manuscript. We further acquired an anatomical scan at the start of each measurement applying a 3D T1-weighted Magnetization Prepared-RApid Gradient Echo (MP-RAGE) sequence (TR=2300 ms; TE=2.94 ms; TI=900 ms; flip angle=9°; voxel size=1×1×1 mm^3^; FOV=320×320mm^2^).

### Neuroimaging study: analysis of behavioral and hormonal data

For the analysis of behavioral data, we first ran a paired t-test comparing mean aggression against in the low- vs high-status condition as we had done in the behavioral study. Additionally, we ran linear mixed-effects models on participants’ aggressive behavior, decision latencies, and reaction times. We specifically tested for effects of opponent status (high or low), trial number (1 to 45), and outcome of the previous trial (won or lost). These three variables were defined as fixed effects whereas subject was defined as random effect. We included by-participant slopes for all fixed effects and random intercepts for all predictors where possible, adhering to standard guidelines for linear mixed-effects modeling (Barr *et al.*, 2013). Models with random slopes for outcome failed to converge, just as those with random intercepts for status and outcome. Therefore, all models included random intercepts for subject and trial as well as random by-participant slopes for status and trial. We performed these analyses using the lmerTest package (Kuznetsova *et al.*, 2017) in R (version 3.6.1) running on R Studio (version 1.1.423).

For testosterone and cortisol, we compared the pre- and post-scan log-transformed values using paired t-tests in order to inspect for state changes in hormonal concentrations. Two pre-scan testosterone samples had to be discarded because the amount of saliva did not suffice for reliable quantification. We then tested whether mean aggression against the high- minus the low-status opponent was associated with baseline testosterone, cortisol, the testosterone-to-cortisol ratio, and/or pre-post changes in either hormone using Pearson correlation coefficients. For paired t-tests, we report Cohen’s d_rm_, which accounts for the correlation between repeated measurements (Lakens, 2013). For correlation and regression coefficients, we converted the corresponding r and t-values to Cohen’s d using the *psych* package’s functions t2d and r2d respectively (Revelle, 2017). We used ggplot2 to plot results (Wickham, 2016).

### Neuroimaging study: analysis of fMRI data

We performed the pre-processing and analysis of neuroimaging data using Statistical Parametric Mapping 12 (SPM12; https://www.fil.ion.ucl.ac.uk/spm/) running on Matlab 2019b. We used a standard pre-processing pipeline that involved, in this order, slice-timing correction to the four middle slices (i.e. those acquired at 485ms) with a 4^th^ degree spline interpolation, realignment to the first functional volume, coregistration of mean functional and anatomical images, segmentation of the anatomical images based on default tissue probability maps, normalization to the Montreal Neurological Institute (MNI) template and smoothing with an 8mm full-width at half-maximum Gaussian kernel. Additionally, we performed distortion correction using the FieldMap toolbox, except in seven participants for which this procedure failed or worsened image quality. We excluded one imaging run in three subjects due to excessive head movements (>3mm or 3° in any direction).

First-level models comprised two regressors for the decision phase (high- or low-status opponent; 4s) and four for the outcome phase (won or lost vs high- or low-status; 4s). We also included the absolute distance in status between subject and opponent in percentage as a covariate in the decision phase to control for momentary fluctuations in status. As regressors of no interest, we modelled the warning sign, target, and motor responses in the reaction time task as well as the sound at the end of the outcome phase as delta functions with null duration. We further included the six movement parameters estimated from realignment (x, y, z, pitch, roll, and yaw). We convolved all regressors (except for motion parameters) with the canonical hemodynamic response function, implemented a high-pass filter with a 128s cut-off, and applied the standard SPM autoregressive model AR(1) to control for autocorrelation. In order to inspect for trial-wise changes in activity associated with aggression, we additionally defined identical first-level models that included trial-wise mean-centered punishment selections as parametric modulator.

We first performed multi-voxel pattern analyses (MVPA) to uncover local activation patterns that most differentiated between the two opponents in the decision phase. We used the Decoding Toolbox (Hebart *et al.*, 2015) running on Matlab 2019b to implement a linear support vector machine algorithm that classifies multivariate activation patterns for each condition and run using a leave-one-out cross validation procedure on the corresponding beta images. We ran such an analysis within each of eight regions-of-interest (ROIs) putatively involved in the relational processing of social status (Zerubavel *et al.*, 2015; Qu *et al.*, 2017; Schafer and Schiller, 2018). We defined these ROIs as bilateral anatomical masks extracted from the Automatic Anatomical Labelling atlas (Maldjian *et al.*, 2003): the amygdala (merge of left and right “Amygdala” masks), anterior cingulate cortex (“Cingulum_Ant” mask), dorsomedial prefrontal cortex (“Frontal_Sup_Medial” mask), hippocampus (merge of left and right “Hippocampus” masks), precuneus (“Precuneus” mask), striatum (merge of “Caudate”, “Putamen”, and “Pallidum” masks), temporal-parietal junction (merge of “Temporal_Sup” and “SupraMarginal” masks), and ventromedial prefrontal cortex (merge of left and right “Rectus” and “Frontal_Med_Orb” masks). We resampled the masks to match the voxel size of the functional data (3mm). For each of these regions, we extracted the participant-wise area under the curve (AUC) minus chance as a classification performance measure. Here, AUC expresses the total surface below the receiver operating characteristic curve, which is obtained by plotting the cumulative true positive rates against the cumulative false positive rates. Hence, higher AUC values indicate a greater capacity to classify between categories. While standard accuracy measures treat all classifications equally, AUC also takes into account the “confidence” (i.e. the distance to the decision boundary) that the classifier has regarding the membership of specific items (Hebart *et al.*, 2015). In order to assess statistically which ROIs distinguished between conditions, we computed the bootstrapped 95% bias-corrected accelerated confidence intervals (BCa CI) confidence intervals of the average AUC for each ROI. ROIs whose CI did not include zero were considered to classify between the high- and low-status rival at an above-chance level. In order to more precisely localize the observed effects, we performed a post-hoc searchlight analysis (9-voxel radius sphere) within each ROI showing above-chance classification. These analyses were thresholded at p<.05, k>50 uncorrected.

We subsequently sought for associations between neural reactivity to status signals and aggression on a within-subject basis. We did so by comparing the parametric modulator for punishment selections in the low- versus the high-status condition using a paired t-test. This analysis tests for brain areas showing transient increases in activity as a function of participants’ trial-wise aggression levels against one opponent relative to the other. Five participants had to be excluded from this analysis due to lack of variability in behavior (i.e. they chose the same punishment in all trials). Given that we had no a priori hypotheses for these analyses, we applied a threshold of p<.001 at the voxel level with a p<.05 family-wise error (FWE) correction at the cluster level.

In the outcome phase, we tested whether wins and losses were differentially processed depending on status using a flexible factorial analysis. First, we tested the main effects of won vs lost in order to reproduce the VS and vmPFC activity commonly observed during competitive victories. Because subjects lost more often against one of the two opponents, status and outcome were not orthogonal. We therefore did not inspect the main effect of status. Rather, we directly tested the interaction between the two factors, namely the contrast ***[won high + lost low] > [lost high + won low]*** and its opposite using paired t-tests (see Table S1 in Supplementary Material), and post-hoc plotted the resulting parameter estimates. For these contrasts we also set a voxel-level p<.001 threshold with a cluster-wise pFWE<.05 correction, as the effect of competitive outcomes on the targeted valuation areas is usually detectable in whole-brain analyses (Votinov *et al.*, 2015; Ligneul *et al.*, 2016).

Finally, we probed whether interindividual differences in status-based aggression were associated with the neural processing of status signals. We used MarsBaR (http://marsbar.sourceforge.net/) to extract subject-wise parameter estimates from ROIs showing above-chance classification in the MVPA analyses (whole ROI) and from clusters showing significant activity in the outcome by status interaction (6 mm sphere around peak coordinate). We computed the correlation coefficients between activity in each ROI/cluster and mean aggression against the high-minus the low-status opponent. We also computed the 95% bootstrapped confidence intervals for the correlation coefficients of significant associations (p<.05) in order to assess their robustness and to protect against the influence of outlying observations. We performed all bootstrapping analyses using the *bootstrap* package (version 2019.5) in R (version 3.6.1) running on R Studio (version 1.1.423). We used ggplot2 to depict all neuroimaging results (Wickham, 2016). Behavioral and ROI data along with analysis scripts for this study are freely available via the Open Science Foundation (https://osf.io/2jvx4/). Raw and pre-processed neuroimaging data are available upon request.

### Neuroimaging study: behavioral results

Participants were 47 healthy young men (age=24 ± 3.8 [mean ± standard deviation]) who believed to be playing against two other participants. These were actually confederates and the task was again preprogramed. Here, the status manipulation was also successful, as subjects were aware of winning less often against the stronger than against the weaker rival (t_46_=10.05, d=2.36, p<.001). Subjects overestimated the percent of trials won against both the better (41.1% ± 1.6% [mean ± standard error] vs 33%, t_46_=4.74, d=.69, p<.001) and worse opponents (71.9% ± 1.7% vs 66%, t_46_=3.40, d=.49, p=.001). Again, they did not perceive the stronger opponent to be more aggressive (t_46_=1.26, d=.27, p=.213). As in the pilot study, subjects selected higher average punishments against when they were in a low- than in a high-status position (t_46_=3.57, d=.57, p<.001; Fig. 2c). We thus calculated status-dependent aggression as the difference in average punishment selections in each condition (low- minus high-status) to be used in later correlation analyses with neural and hormonal data.

We additionally tested whether subjects’ behavior changed over time using linear mixed-effects models (see Methods). Participants selected louder sound blasts over time when they were in a low-status position, whereas their punishment selections remained unchanged in a high-status one (status x trial interaction: β=0.125, t_4062_=2.70, p=.006; Fig. 2c; Table 2a). Specifically, they evinced an increase of around .7 points (≈8.75% of the scale) in the last (4.09 ± .08 [mean ± standard error]) compared to the first five trials (3.39 ± .21) when low in status. In contrast, their average aggression levels remained virtually unchanged as high-status players (first five trials: 3.47 ± .14; last five trials: 3.49 ± .07). Thus, participants selected stronger punishments in a low rank even when provocation remained constant.

**Table 2:** Results of linear-mixed effects models in the neuroimaging study (n=47)

| *a) Dependent variable: punishment selections* |  |  |  |  |
| --- | --- | --- | --- | --- |
| *Predictor* | $\beta$ | SE | t | p |
| **Status** | **0.488** | **0.146** | **3.347** | **.002** |
| **Trial** | **0.141** | **0.059** | **2.383** | **.020** |
| Outcome | 0.005 | 0.035 | 0.132 | .895 |
| **Status \* Trial** | **0.125** | **0.046** | **2.708** | **.007** |
| Status * Outcome | 0.014 | 0.048 | 0.288 | .773 |
| Trial * Outcome | 0.049 | 0.035 | 1.394 | .164 |
| Status * Trial * Outcome | 0.041 | 0.048 | 0.85 | .395 |
| *b) Dependent variable: response latency in decision phase* |  |  |  |  |
| Status | 0.026 | 0.018 | 1.426 | .154 |
| Trial | 0.020 | 0.020 | 0.977 | .331 |
| **Outcome** | **0.032** | **0.013** | **2.346** | **.019** |
| Status * Trial | 0.023 | 0.018 | 1.261 | .207 |
| Status * Outcome | 0.022 | 0.019 | 1.163 | .245 |
| Trial * Outcome | 0.008 | 0.014 | 0.606 | .545 |
| Status * Trial * Outcome | 0.003 | 0.019 | 0.168 | .867 |
| *c) Dependent variable: response latency in reaction time task* |  |  |  |  |
| Status | 0.001 | 0.003 | 0.591 | .554 |
| **Trial** | **0.007** | **0.002** | **2.989** | **.004** |
| Outcome | 0.002 | 0.002 | 1.097 | .273 |
| **Status \* Trial** | **0.008** | **0.002** | **3.258** | **.001** |
| Status * Outcome | 0.002 | 0.003 | 0.656 | .512 |
| Trial * Outcome | 0.002 | 0.002 | 0.821 | .412 |
| Status * Trial * Outcome | 0.002 | 0.003 | 0.737 | .461 |

$\beta$ : regression coefficient, SE=standard error, t: t value, p: p-value. Bold values indicate effects significant at $p < .05$ .

Response latency in the decision phase did not vary as a function of status, time, or their interaction (all p>.154; Table 2b). That is, subjects took similarly long to choose the punishment against either opponent across trials (high-status: 1.32 ± 0.16s; low-status: 1.29 ± 0.12s). Nevertheless, there was a significant main effect of outcome (β=0.032 ± 0.01, t_1073_=2.34, p=.019) such that participants chose about 50ms faster after losing (1.27 ± 0.01ms) than after winning (1.32 ± 0.01s).

In the reaction time task, participants became progressively quicker as low-status players (status-by-trial interaction: β=0.008, t_4111_=3.25, p=.001; Fig. 2d; Table 2c). On average, subjects were approximately 22ms faster in the last five (3.50 ± 0.03s) relative to the first five trials (3.72 ± 0.06s) against when they had a low status, whereas they were comparably quick throughout the task in the high-status condition (first five trials: 3.64 ± 0.12s; last five trials: 3.65 ± 0.06s). This confirms the motivational relevance of competitive status for participants’ performance.

### Neuroimaging study: testosterone and cortisol results

Concerning hormonal concentrations, average testosterone levels increased 1.57% after scanning (t_44_=2.10, d=.36, p=.040; Fig. 2d) whereas mean cortisol decreased by 39.74% (t_46_=4.96, d=.92, p<.001; Fig. 2e). Baseline concentrations of testosterone (r_43_=.08, p=.595) or cortisol (r_45_=−.01, p=.973) were not associated with aggression, nor was the baseline testosterone-to-cortisol ratio (r_43_=.10, p=.497) or the pre-post change in either hormone (testosterone: r_43_=.07, p=.627; cortisol: r_45_=.06, p=.686). Therefore, we did not use endogenous hormone levels for further analyses.

### Neuroimaging study: status-dependent activity patterns during punishment selection

With regards to neuroimaging data, we first tested whether regions-of-interest (ROIs) previously linked to the processing of status signals also encode competitive status during decisions to aggress (Fig. 3a). To that end, we performed multi-voxel pattern analysis (MVPA) in the decision phase (see Methods). This revealed that the striatum and hippocampus distinguished between opponents (Fig. 3b). That is, multivariate activity patterns in these regions could classify at an above-chance level whether subjects were facing the high- or the low-status opponent. Post-hoc searchlight analyses showed that both ventral and dorsal aspects of the left striatum contributed to the effect, whereas in the case of the hippocampus classification was most strongly driven by its right anterior portion (Fig. 3c). We then tested whether a stronger multivariate differentiation between opponents was linked with status-dependent aggression across subjects. To do so, we took classification performance values for the hippocampus and striatum and correlated them with mean aggression against the high- minus low-status opponent. Although we observed no effects for the hippocampus (r=_45_-.03, p=.823), there was an association between striatum activity patterns and aggression against the high- minus low-status rival (r_45_=.29, d=.62, p=.043; Fig. 3d). In other words, subjects showing a more pronounced differentiation between opponents in the striatum showed greater status-contingent aggressive behavior.

**Fig. 3.**
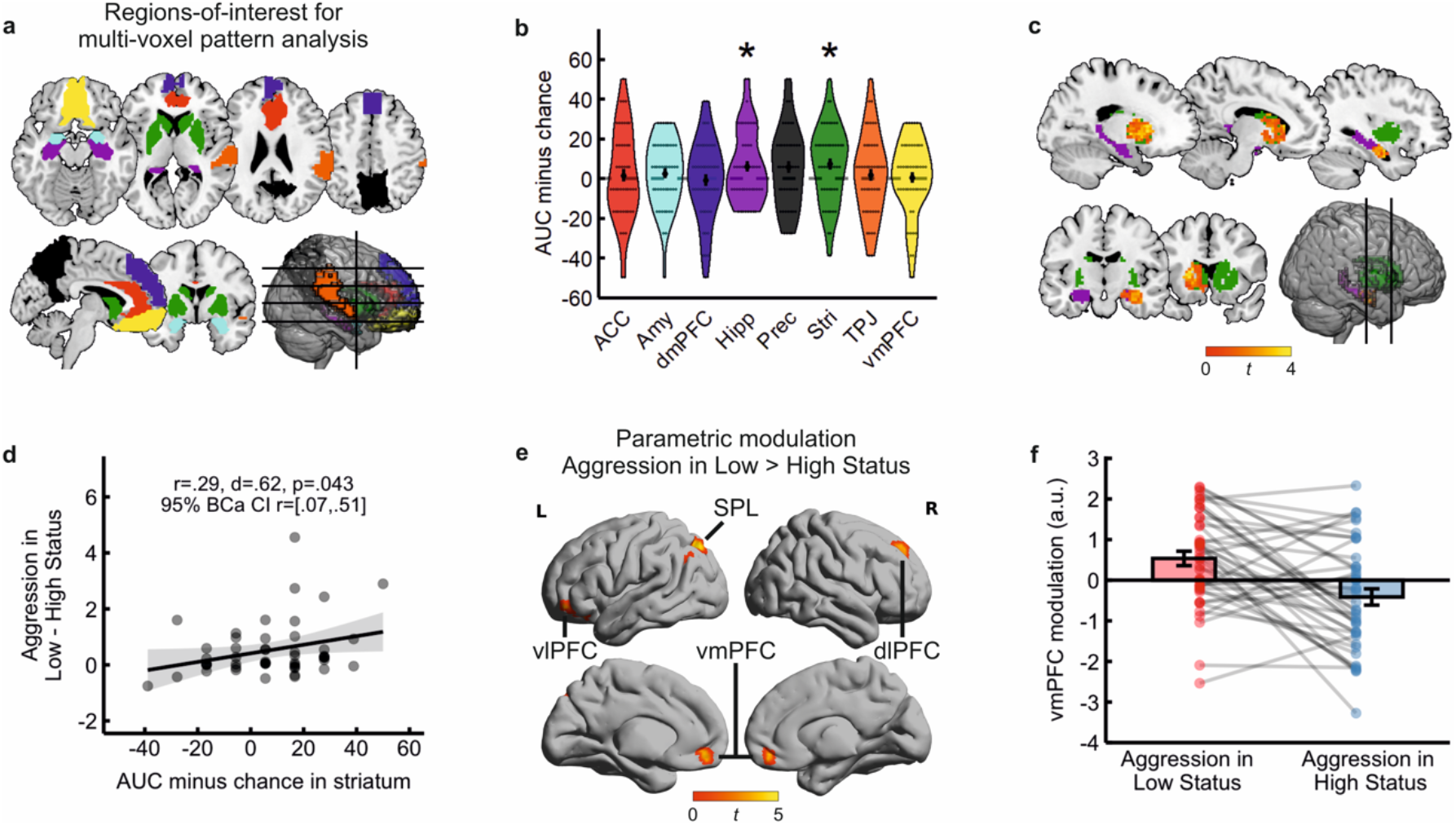
Neural processing of competitive status during punishment selection (decision phase). **a**, regions-of-interest (ROIs) used in multi-voxel pattern analysis (MVPA) to identify multivariate status representations during punishment selection. **b**, classification performance expressed as area under the curve (AUC) minus chance per each ROI. Each dot shows the AUC value per subject, with higher values indicating better classification between the better- and worse-performing rivals. Asterisks denote ROIs with significant above-chance accuracy according to bootstrap resampling. ACC: anterior cingulate cortex; Amy: amygdala; dmPFC: dorsomedial prefrontal cortex; Hipp: hippocampus; Prec: precuneus; Stri: striatum; TPJ: temporal-parietal junction; vmPFC: ventromedial prefrontal cortex. **c**, results of the post-hoc searchlight MVPA analysis within hippocampus and striatum masks (voxel-level p<.05 uncorrected, cluster size k>50). Higher values indicate a greater contribution to the classification. **d**, correlation between classification performance in the striatum and mean aggression against the high-minus low-status opponent. We include best-fit lines and 95% prediction intervals. r=Pearson correlation coefficient; d=Cohen’s d; 95% BCa CI: 95% bias-corrected accelerated confidence intervals. **e**, parametric modulation in the decision phase (voxel-level p<.001, cluster-level p<.05 Family Wise-Error [FWE] corrected, n=42). Higher values indicate more trial-wise activity as a function of aggression in a low- relative to a high-status position. dlPFC: dorsolateral prefrontal cortex; SPL: superior parietal lobe; vlPFC: ventrolateral prefrontal cortex. **f**, parameter estimates resulting from the parametric modulation in the vmPFC. A.u.: arbitrary units.

### Neuroimaging study: trial-by-trial fluctuations in punishment selections

We subsequently inquired whether neural reactivity to status cues influenced punishment selections on a within-subject basis. We hence performed a parametric modulation analysis comparing trial-wise decisions against in each condition (see Methods). As shown in Fig. 3e, greater aggression against in a low- compared to a high-status position was associated with *increased* activity in ventromedial prefrontal cortex (vmPFC; t=4.57, k=59, x=9, y=50, z=−10; Fig. 3b), dorsolateral prefrontal cortex (dlPFC; t=4.11, k=57, x=21, y=44, z=41), ventrolateral prefrontal cortex (vlPFC; t=4.79, k=57, x=−33, y=32, z=−16) and superior parietal lobe (SPL; t=4.80, k=88, x=−24, y=−76, z=47). That is, activity in these regions co-varied with participants’ aggressive behavior in a low-relative to a high-status rank. No regions survived in the opposite contrast (high > low).

### Neuroimaging study: neural processing of competitive outcomes as a function of status

In the outcome phase, we observed widespread activation in the contrast won > lost with peaks in the bilateral ventral striatum (VS; Table S1 in Supplementary Material). This is in line with previous results employing competitive tasks (Votinov *et al.*, 2015; Buades-Rotger, Brunnlieb, *et al.*, 2016). More importantly, we tested the interaction between status and outcome in order to inspect whether victories and defeats were differently processed in the low-compared to the high-status condition. The interaction contrast ***[Won Low + Lost High)] > [Lost High + Won Low]*** revealed activation in left anterior insula (AI), dorsal anterior cingulate cortex (dACC), left dorsolateral prefrontal cortex (dlPFC), left and right temporoparietal junction (TPJ), precuneus, and VS (Fig. 4a-c; see Table S1 in Supplementary Material for complete results). These regions hence reacted more strongly to infrequent, status-incongruent outcomes. However, as becomes apparent from the parameter estimates in the VS and dACC (Fig. 4b-c), the interaction was driven by a stronger effect of the outcome when playing against the high- relative to the low-status opponent. If the effect were merely driven by the infrequency or unpredictability of events, activity should be also increased in lost relative to won trials against the low-status opponent, which was not the case. The opposite contrast, which entailed expectable, status-congruent outcomes, yielded activation in primary visual and inferior temporal cortex as well as a cluster in the brainstem (see Table S1 in the Supplementary Material).

**Fig. 4.**
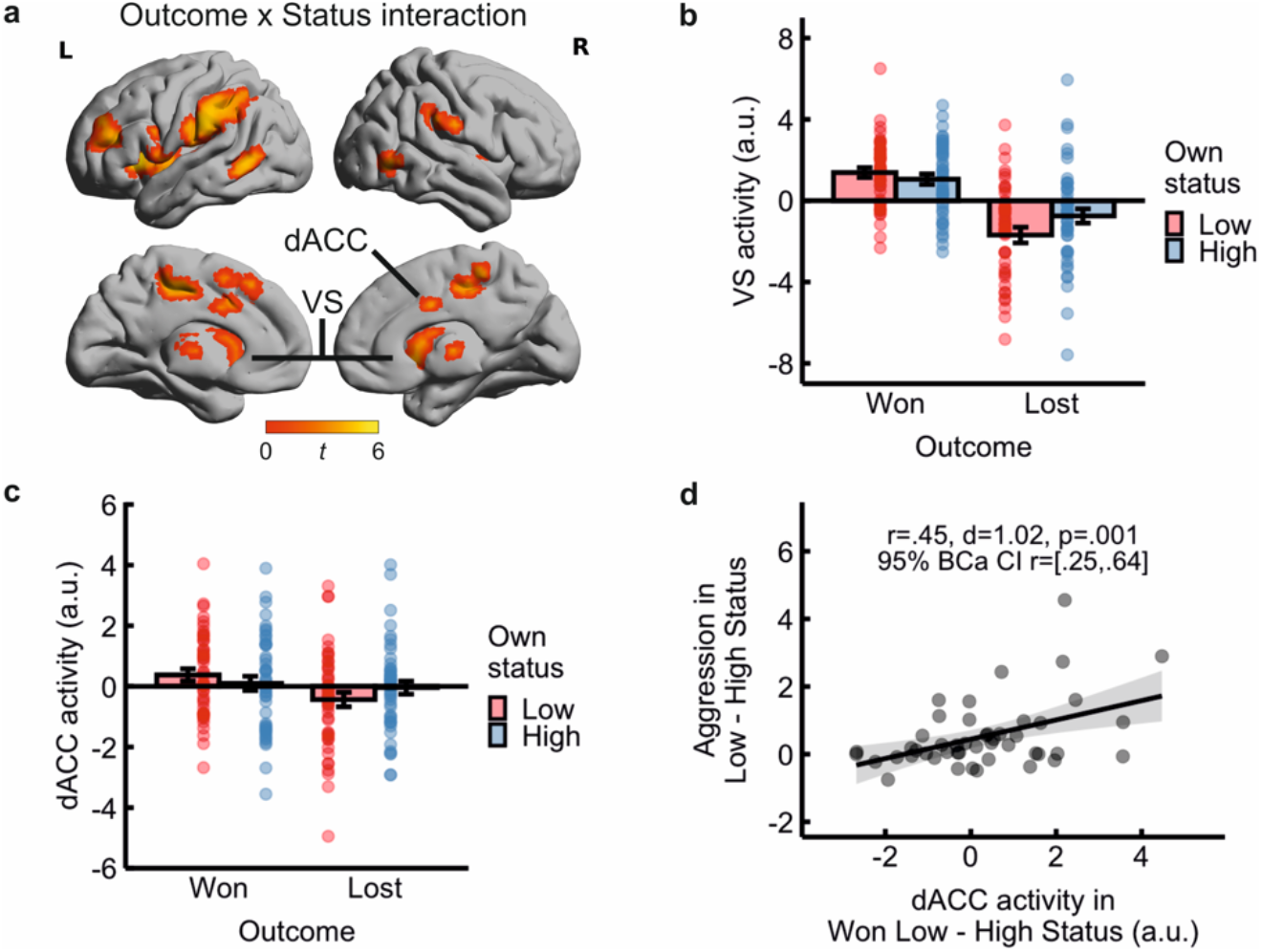
Neural processing of competitive victories and defeats (outcome phase). **a**, regions showing a significant interaction between status and outcome (voxel-level p<.001, cluster-level p<.05 Family Wise-Error [FWE] corrected). **b**, parameter estimates for the interaction effect in the ventral striatum (VS). **c**, parameter estimates for the interaction effect in the dorsal anterior cingulate cortex (dACC). **d**, correlation between dACC reactivity to victories and aggression in the low- minus high-status condition, including best-fit lines and 95% prediction intervals. r=Pearson correlation coefficient; d=Cohen’s d; 95% BCa CI: 95% bootstrapped bias-corrected accelerated confidence intervals (BCa CI).

We next probed whether the neural processing of victories and defeats was linked with status-dependent aggressive behavior. We therefore extracted parameter estimates from clusters resulting from the ***[Won High + Lost Low)] > [Lost High + Won Low]*** contrast (dACC, AI, dlPFC, left and right TPJ, precuneus, and VS) and correlated them with the status effect on aggression. Specifically, we extracted the high- minus low-status difference separately for victories and defeats. Because we performed 14 correlations (2 values for each of 7 ROIs), we applied a Bonferroni correction to control for multiple comparisons (.05/14=.003). Differential reactivity to victories in the dACC was significantly associated with aggression against the high- minus the low-status opponent (r_45_=.45, d=1.02, p=.001; Fig. 4d). There was no correlation for losses (r_45_=.12, p=.394). Activity in the rest of ROIs (AI, dlPFC, left and right TPJ, and VS) was not associated with status-dependent aggression at a corrected level (all p>.050; see Table S2 in Supplementary material).

## Discussion

### Low competitive status is associated with aggression

Aggressiveness is often taken as a requisite for competitive success (Chow *et al.*, 2009; Trebicky *et al.*, 2013; Waasdorp *et al.*, 2013; Laustsen and Petersen, 2017; Cheng, 2020; Weick, 2020). However, better contenders can rely on their skill, whereas less successful ones may use aggression to hamper rivals’ progress (Kirker *et al.*, 2000; Coulomb-Cabagno and Rascle, 2006). Moreover, the frustration and loss of control entailed in being repeatedly defeated should facilitate rather than suppress aggression in low-status contestants (Przybylski *et al.*, 2014; Yu *et al.*, 2014; Oxford *et al.*, 2017). In a correlational analysis and two laboratory experiments we found that low competitive status, measured as competitors’ relative rank in a hierarchy, was linked with more frequent and intense aggressive behavior. Our results replicate and extend previous investigations showing that fouls or penalties are associated with worse overall team performance in American football (Hauge, 2012), ice hockey (Coates *et al.*, 2012), soccer (Jewell, 2012), and basketball (Berri and Rodenberg, 2012). Unlike these studies, however, our measure of aggression (fouls or penalties committed relative to received) captures the reciprocal nature of aggressive play and thus provides clearer evidence on the status-aggression relationship.

The findings from our two experimental studies further show that the link between low status and aggression holds in competitions between individuals (i.e. not only between teams) and when aggressive behavior does not directly offer a clear competitive advantage. In addition, in our neuroimaging experiment subjects became quicker and selected louder sound blasts over time against in a low-status position, even when the opponents’ hostile intent remained constant over time. This finding further demonstrates that aggression increases as competitive hierarchies emerge and is not a mere correlate of low performance. Our results also imply that the relationship between competitive status and aggression is not limited to situations involving face-to-face competition, but also applies to virtual, indirect contests. In line with this assumption, an online bargaining study showed that individuals attribute more hostile intentions to higher-status opponents (Saalfeld *et al.*, 2018). Competitive status thus appears to be a relevant trigger for aggressive behavior in both direct and virtual interactions.

Our main finding that participants chose higher punishments when placed in a low rank is however not entirely expectable. After all, participants might have instead displayed the opposite pattern, i.e. punishing a weaker rival to assert their dominance while being less aggressive against a stronger opponent as a conciliatory strategy. Indeed, some studies have shown that participants accept more unfair offers in experimentally-induced low status (Blue *et al.*, 2016; Hu *et al.*, 2016). Nevertheless, in these investigations behaving submissively yielded a direct economic benefit. What could explain the results of the present study, in which there was no money at stake? One possibility is that subjects were more willing to incur the risk of behaving aggressively when their status was low, as the prospect of losing increases risk-seeking (Tversky and Kahneman, 1981; Ruggeri *et al.*, 2020). Conversely, being in a high-status position induces risk aversion, so that subjects might avoid punishing lower-status individuals due to e.g. reputational concerns (Dreber *et al.*, 2008). Put otherwise, subjects had less to lose in a low-status position, which facilitates risk-taking. In addition, inducing a low-status mindset threatens self-worth and increases hostility (Davis and Reyna, 2015), which might have further exacerbated status-dependent aggression in the present study. Future studies should disambiguate these putative mechanisms underlying status-based aggression.

### Changes in testosterone and cortisol concentrations

We also observed a post-task increase in testosterone concentrations. This surge occurred instead of the decline that could be expected due to the hormone’s circadian rhythm (Keevil *et al.*, 2013; Buades-Rotger, Engelke, *et al.*, 2016). In men, testosterone tends to rise after winning (Geniole *et al.*, 2017) and when achieving a higher status in skill-based hierarchies (Cheng *et al.*, 2018). Although subjects in the present study only won half of the trials on average, they did clearly defeat one opponent and overestimated the percentage of trials won against both rivals. Indeed, elevations in testosterone levels have been linked to more positive self-appraisals of competitive performance (Casto *et al.*, 2017). However, our data do not allow to determine whether the changes we observed here were purely driven by competition itself, the resulting outcomes, or other factors. Cortisol concentrations on the other hand showed a steep decline after the task. This effect, which we have previously observed (Buades-Rotger, Engelke, *et al.*, 2016), is likely due to anticipatory arousal, circadian decline, and/or relaxation in the scanner; the latter might have been heightened by the post-task resting-state measurement. Nonetheless, contrary to prior studies (Carré and Olmstead, 2015), neither endogenous testosterone nor cortisol were linked with status-dependent aggressive behavior. Our results are also in partial disagreement with the finding that competition-related surges in testosterone facilitate further antagonistic behavior (Mehta and Josephs, 2006; Carré *et al.*, 2013), although in the present study aggression was measured *during* competition rather than after it. Many experimental studies on the topic subject participants to rigged competitions in which they are clear winners or losers (Geniole *et al.*, 2017). In contrast, the task employed here entailed a sequence of wins and losses, which might have had mutually cancelling effects on testosterone dynamics and subsequent aggressive impulses. Furthermore, testosterone increases are stronger when measured more than 10 minutes before competition (Geniole *et al.*, 2017). Our design might have hence missed out on the preparatory testosterone rush posited to foster competitive aggression in men (Zilioli and Bird, 2017; Geniole and Carré, 2018). Recent meta-analytic evidence suggests that the association between testosterone and aggression in humans is small (Geniole *et al.*, 2020). Testosterone-by-cortisol interactions yield even lower effect sizes across studies, with highly variable estimates and some evidence for publication bias (Dekkers *et al.*, 2019). Large, pre-registered investigations (Brannon *et al.*, 2019) are necessary to reach conclusive evidence on the role of testosterone in status-based aggression.

### Neural representations of competitive status during punishment selection

At the neural level, we identified a number of brain regions implicated in the processing of competitive hierarchies and characterized their relevance for status-dependent aggressive behavior. Multivoxel pattern analyses revealed that the right anterior hippocampus distinguished between the better and worse rivals during punishment selection. A growing number of studies indicate that the role of the human anterior hippocampus is not limited to mapping events in time or items in space. Rather, this structure has been suggested to encode abstract relationships between entities along several dimensions extracted from statistical regularities in the environment, i.e. relational knowledge (Garvert *et al.*, 2017; Park *et al.*, 2020). Similar mechanisms have been documented in the social domain, so that the anterior hippocampus tracks changes in an individual’s power and affiliation relative to oneself (Tavares *et al.*, 2015; Kumaran *et al.*, 2016). Our results therefore provide converging evidence that this region, analogous to the ventral hippocampus in rodents (Fanselow and Dong, 2010), encodes status relationships and can thereby inform social decision-making.

The left ventral and dorsal striatum also displayed multivariate signals distinguishing the high- and low-status rivals. Furthermore, participants with a stronger differentiation between opponents in the striatum showed more status-dependent aggression. Paralleling our findings, striatum neurons in the rhesus macaque brain respond differentially to social cues from dominant and submissive monkeys, which concurs with a preference to watch faces from higher-status individuals (Klein and Platt, 2013). The present results thus indicate that the striatum is not only sensitive to status signals (Zink *et al.*, 2008; Zerubavel *et al.*, 2015) and competitive outcomes (Qu *et al.*, 2017), but also differentiates between individuals differing in status during decisions to aggress. Our findings are in line with the hypothesized role of the striatum in retaliatory aggression (Chester, 2017; Bertsch *et al.*, 2020), and concur with meta-analytic findings demonstrating increased striatal activity when individuals deliver harsher punishments to unfair co-players (Gabay *et al.*, 2014). Therefore, our data bridge animal and human research in showing that the hippocampus and striatum are involved in the relational processing of social dominance signals. The present findings additionally suggest that these mechanisms are at play during competitive interactions and might contribute to the adoption of aggressive strategies.

### Trial-wise covariation between brain activity and status-dependent aggression

The ventromedial prefrontal cortex (vmPFC) evinced a status-dependent covariation with participants’ momentary changes in aggressive behavior. Put another way, this region integrated status information in decisions to aggress on a trial-by-trial level. A number of volumetric and functional studies indicate that the vmPFC is crucial for the control of aggressive impulses (Beyer *et al.*, 2015; Gilam *et al.*, 2015; Chester *et al.*, 2017), presumably by downregulating amygdala activity (Coccaro *et al.*, 2011; Motzkin *et al.*, 2015). There are however contradictory findings showing that vmPFC damage is linked with *increased* cooperative behavior (Wills *et al.*, 2018), or that vmPFC activation *positively* predicts aggressive decisions (Buades-Rotger *et al.*, 2017; Repple *et al.*, 2017). Here, the status-contingent, aggression-related elevations in vmPFC activity went along with those of other regions putatively involved in emotion regulation and social decision-making such as the dorsolateral or ventrolateral prefrontal cortex (Morawetz *et al.*, 2015; Hackel *et al.*, 2020). In contrast, there were no positive associations between brain structures assumed to generate aggressive urges (e.g. amygdala, periaqueductal grey) (Coccaro *et al.*, 2011; Yu *et al.*, 2014) and punishment selections in the task. Therefore, we speculate that the observed prefrontal and parietal activity might correspond to deliberate, intentional aggressive decisions. This assumption is supported by the finding that lost trials led to subsequently faster decisions but did not influence punishment selections, which were strongly driven by competitive status. The relatively long and invariant response times during punishment selection (i.e. around 1300ms throughout the task) are also in consonance with this interpretation. Indeed, there is evidence that willingness to influence the rival’s performance accounts for some variation in average punishment selections in this task (Chester and Lasko, 2019). Nonetheless, as we did not measure participants’ motives, we cannot determine to which extent the intent behind louder sound blasts was mostly hostile or instrumental in nature.

### Status-dependent reactivity to competitive outcomes

Finally, there was enhanced activation for status-incongruent outcomes in the ventral striatum, anterior insula, dorsal anterior cingulate cortex (dACC), dorsolateral prefrontal cortex, temporoparietal junction (TPJ) and precuneus. More specifically, these regions showed a stronger differentiation between victories and defeats when participants were in a low- relative to a high-status position. Our results resemble those from a previous study in which subjects competed against rivals varying in status (Zink *et al.*, 2008), and further implicate the VS in competitive hierarchy learning (Ligneul *et al.*, 2016). The present findings are also consistent with the observation that brain areas involved in valuation (e.g. VS) and social cognition (e.g. TPJ, precuneus) display differential sensitivity to a person’s position in a hierarchy (Zerubavel *et al.*, 2015). In addition, dACC reactivity to victories in a low as compared to a high rank predicted greater aggression in the former relative to the latter. That is, the dACC response to status-enhancing wins, as compared to status-asserting ones, was correlated with greater status-dependent aggression. The local maximum observed here was located in the most posterior aspect of the dACC and lied in the near vicinity (i.e. within 5 mm in the sagittal and axial planes) of a meta-analytic peak for vicarious pain (Lamm *et al.*, 2011). Concordantly, dACC activity has been linked with retaliatory behavior (Krämer *et al.*, 2007; Gabay *et al.*, 2014; Beyer *et al.*, 2015). In rodents, this brain region was reported to encode competitive effort, i.e. the net value of a reward when competition is required to attain it (Hillman and Bilkey, 2012). It is nonetheless puzzling that the dACC failed to show a clear status distinction in the decision phase. This insinuates that, in the context of competitive aggression, the dACC is more responsive to *changes* in status than to status cues per se. In consonance with this observation, the cingulate gyrus has been postulated to track moment-to-moment fluctuations in others’ motivation during social interactions (Apps *et al.*, 2016) as well as in their rank and power within a hierarchy (Kumaran *et al.*, 2016). Taken together, these results tentatively suggest that status-based aggression might be particularly pronounced in persons for whom status-enhancing victories are more salient.

### Limitations

There are at least three key limitations to our experimental studies. First, both samples were small and restricted to healthy young men from whom we did not collect ethnicity data. The generalizability of our findings is thus modest. Second, as commented above, we did not address participants’ strategies or intent in the task. Although reaction time and neuroimaging results may provide some indirect evidence in this regard, we cannot ultimately ascertain whether subjects’ punishment selections were predominantly driven by spiteful or instrumental motives, nor whether they were influenced by frustration, threatened self-esteem, or other affective processes. Third, participants did not face a same-status rival, which would have constituted an ideal control condition and allowed to answer additional research questions. It should be noted that confederates (five male students aged 20-30) were not the same in all measurements, but this is unlikely to be a major source of variability because participants did not know the identity of the other players during the game.

The correlational sports data is limited in its correlational nature and in that we did not separately consider direct encounters between low- and high-status teams. Single-match metrics that model the distance in rank between opposing teams would more closely parallel the behavioral and neuroimaging studies. Finally, sports data only included men for consistency with the experimental studies and this also curtails the extrapolation of the obtained results to other competitions.

## Conclusions

In sum, we found that low competitive status consistently evokes aggression in healthy young men. Our findings refute the notion that successful competitors are the more aggressive ones. Rather, our results indicate that those in the lower echelons of competitive hierarchies use aggression more frequently and intensely over the course of successive encounters. Our experimental data shows that this holds even for situations in which aggression does not directly serve any instrumental function. Although endogenous testosterone levels increased after a competitive interaction, there were no links between this hormone and aggression, neither alone nor in interaction with cortisol. The latter finding questions a strong impact of testosterone on status-driven aggression, in line with recent meta-analytic evidence (Dekkers *et al.*, 2019; Geniole *et al.*, 2019). On a neural level, both anterior hippocampus and striatum displayed multivariate representations of competitive status. Interindividual differences in status-dependent aggressive behavior could be predicted by a stronger differentiation between opponents in the striatum and by neural reactivity to status-enhancing victories in the dACC. On a trial-by-trial, within-subject basis, the vmPFC together with other prefrontal and parietal regions favored status-based aggression. Our study thus reveals a consistent association between low competitive status and aggression, and points toward potential neural mechanisms underlying this relationship.

## Acknowledgements

This study was funded by the German Science Foundation (grant number KR3691/5-1). We are grateful to Susanne Schellbach and Mourad Zoubir for their help with the data acquisition and to our subjects and confederates for their participation. The authors report no conflicts of interest.

