## Supplementary Material for "Low competitive status elicits aggression in healthy young men: behavioral and neural evidence"

**Table S1**. Whole-brain results in the outcome phase

| *Won > Lost*[a] | k | t | x y z |
|---|---|---|---|
| Ventral striatum | 859 | 15.26 | 18, 8, -7 |
| | | 11.28 | 27, -10, 8 |
| | | 9.81 | 21, -1, 14 |
| Angular gyrus | 1384 | 13.85 | -42, -52, 35 |
| | | 13.72 | -42, -58, 44 |
| | | 12.28 | -39, -37, 47 |
| Ventral striatum | 812 | 13.44 | -15, 11, -4 |
| | | 11.22 | -24, -10, 14 |
| | | 10.99 | -21, -4, 2 |
| Dorsolateral prefrontal cortex | 1324 | 12.79 | -39, 20, 44 |
| | | 12.64 | -33, 14, 50 |
| | | 11.70 | -27, 8, 59 |
| Medial temporal gyrus | 398 | 12.77 | -57, -34 -10 |
| Cerebellum | 1830 | 12.46 | 30, -67 -31 |
| | | 11.74 | 3, -55 -25 |
| | | 11.27 | 0, -52 -16 |
| Ventrolateral prefrontal cortex | 198 | 11.62 | -33, 50, -7 |
| | | 11.48 | -39, 53, 5 |
| Supramarginal gyrus | 930 | 11.42 | 45, -28, 44 |
| | | 11.32 | 45, -55, 32 |
| | | 11.07 | 48, -64, 5 |
| Precuneus | 128 | 10.58 | -3, -58, 41 |
| Dorsolateral prefrontal cortex | 589 | 10.38 | 36, 14, 47 |
| | | 10.14 | 27, 20, 50 |
| | | 10.10 | 24, -1, 53 |
| Ventrolateral prefrontal cortex | 141 | 9.74 | 42, 44 -13 |
| | | 9.34 | 45, 35 -13 |
| | | 9.17 | 30, 50, -7 |
| Middle temporal gyrus | 261 | 9.63 | 63 -31 -13 |
| | | 9.27 | 63, -4 -25 |
| | | 8.97 | 51, -46, -7 |
| Precentral gyrus | 30 | 9.39 | 12, -28, 62 |
| *Won < Lost*[a] | | | |
| Superior temporal gyrus | 75 | 12.07 | -39, -34, 11 |
| Superior temporal gyrus | 47 | 10.39 | 48, -28, 11 |
| *(Won High + Lost Low) > (Lost High + Won Low)*[b] | | | |
| Anterior insula | 1132 | 5.81 | -33, 23, 2 |
| Thalamus | | 5.37 | -9 -13, 11 |
| Caudate | | 5.21 | -12, 11, 11 |
| Dorsolateral prefrontal cortex | 240 | 4.96 | -36, 41, 20 |
| Supramarginal gyrus | 497 | 4.88 | -66, -25, 29 |
| | | 4.84 | -63, -31, 35 |
| | | 4.44 | -54, -43, 38 |
| Middle temporal gyrus | 104 | 4.76 | -60, -58, 2 |
| Dorsomedial prefrontal cortex | 62 | 4.49 | -6, 26, 50 |
| | | 3.51 | -6, 17, 56 |
| Posterior cingulate cortex / Precuneus | 240 | 4.41 | -9, -31, 47 |
| | | 4.26 | -6, -22, 47 |
| | | 3.73 | 9, -19, 47 |
| Middle temporal gyrus | 68 | 4.21 | 51, -61, 2 |
| | | 3.96 | 60, -61, -1 |
| Dorsal anterior cingulate cortex | 80 | 3.90 | -9, 20, 26 |
| | | 3.84 | -6, 14, 35 |
| | | 3.74 | 3, 5, 35 |
| Supramarginal gyrus | 61 | 3.73 | 63 -19, 26 |
| | | 3.44 | 54 -19, 29 |
| | | 3.37 | 66 -34, 35 |

| | | | |
|---|---|---|---|
| *(Won High + Lost Low) < (Lost High + Won Low)*[b] | | | |
| Lingual gyrus | 4053 | 9.09 | 18, -79, -10 |
| | | 8.43 | -30, -79, -16 |
| | | 7.64 | -12, -88, -10 |
| Brainstem | 104 | 5.57 | 3, -28, -52 |
| | | 3.96 | -15, -31, -52 |
| | | 3.49 | -6, -28, -55 |

k=cluster size, t=peak t value, x y z: peak coordinates in Montreal Neurological Institute space. [a]t>8 (equivalent to a Family-Wise Error [FWE] correction of p<.000000005), k>20. [b]voxel-level p<.001, cluster-level p<.05 FWE.

**Table S2**. Correlations between brain activity in the outcome phase and the status effect on punishment selection (mean punishment selection against the high- minus low-status opponent)

| *Won High > Won Low* | *r* | *p* |
|---|---|---|
| **Dorsal anterior cingulate cortex** | **.45** | **.001** |
| Anterior insula | .25 | .087 |
| Dorsolateral prefrontal cortex | .07 | .632 |
| Left temporoparietal junction | .23 | .107 |
| Right temporoparietal junction | .26 | .075 |
| Precuneus | .28 | .053 |
| Ventral striatum | .28 | .050 |
| *Lost High > Lost Low* | | |
| Dorsal anterior cingulate cortex | .12 | .394 |
| Anterior insula | .002 | .988 |
| Dorsolateral prefrontal cortex | .10 | .497 |
| Left temporoparietal junction | .13 | .366 |
| Right temporoparietal junction | .09 | .534 |
| Precuneus | .07 | .633 |
| Ventral striatum | -.13 | 359 |

r=correlation coefficient, p=p value. Bold values flag regions showing a significant association with the status effect at a Bonferroni-corrected threshold of p<.003.
